# Field conditions greatly modify a major growth-defense tradeoff in *Arabidopsis thaliana*

**DOI:** 10.1101/2024.08.15.607984

**Authors:** Derek S. Lundberg, Sonja Kersten, Ezgi Mehmetoğlu, Pratchaya Pramoj Na Ayutthaya, Wangsheng Zhu, Karin Poersch, Wei Yuan, Sophia Swartz, David Müller, Ilja Bezrukov, HARVEST TEAM, Detlef Weigel

## Abstract

When plants defend themselves from pathogens, this often comes with a trade-off: the same genes that protect a plant from disease can also reduce its growth and fecundity in the absence of pathogens. One protein implicated in a major growth-defense trade-off in the plant *Arabidopsis thaliana* is ACCELERATED CELL DEATH 6 (ACD6), an ion channel that modulates salicylic acid (SA) synthesis to potentiate a wide range of defenses. Wild plant populations maintain significant functional variation in the *ACD6* gene, with some alleles making the protein hyperactive. In the greenhouse, plants with hyperactive *ACD6* alleles are resistant to diverse pathogens, yet are of smaller stature, their leaves senesce earlier, and they set fewer seeds. We hypothesized that such hyperactive alleles would not only affect the growth of microbial pathogens, but also more generally leaf microbiome assembly in the wild. To test this, we grew plants with hyperactive, standard, and defective *ACD6* alleles in the same field-collected soil, both in climate-controlled conditions and outdoors. We surveyed visual phenotypes, gene expression, hormone levels, seed production, and the microbiome in each environment. To our surprise, we discovered that mature field plants, in stark contrast to greenhouse plants, were unaffected by their *ACD6* genotype, suggesting that additional abiotic and/or microbial signals present outdoors – but not in the greenhouse – greatly modulate *ACD6* activity.

## Introduction

The immune system protects plants from pests and pathogens by detecting foreign molecules [1,2] or physical damage [3] and triggering signaling cascades that greatly increase latent defenses. This includes the production of reactive oxygen and other defense molecules that neutralize or mitigate biotic threats, but that at the same time can also damage the host [2,4]. In the plant *Arabidopsis thaliana*, the membrane-bound ankyrin-repeat protein ACCELERATED CELL DEATH 6 (ACD6) plays an important role in coordinating several immune responses, and related ankyrin-repeat proteins have been implicated in defense in other species including maize [5] and wheat [6]. ACD6 associates with other defense proteins including the plant surface receptor FLS2 that detects bacterial flagellin [7–9], and it acts as an ion channel for calcium [9]. *ACD6* expression, protein accumulation and activity are modulated by the plant defense hormone salicylic acid (SA), and ACD6 in turn regulates SA levels in a positive feedback loop [7].

Similar to many other immune genes, there is substantial structural and sequence diversity in the *ACD6* genomic region across accessions of *A. thaliana* [10–13]. These have a wide range of activities, either on their own or as heteroallelic combinations, and their activity can be further modulated by second-site genetic variants [9–13]. The class of alleles first found in the accession Est-1 (*ACD6*-Est-1) appears hyperactive when compared to the standard allele in the reference accession Col-0 [10]. Relative to standard alleles, *ACD6*-Est-1 increases salicylic acid (SA) accumulation and *PR1* expression, enhances resistance to bacterial, fungal, oomycete pathogens, and likely even sucking insects [10]. These defense benefits are, however, counterbalanced by slower leaf initiation rate, smaller final size, late-onset necrosis in fully expanded leaves, and lower fecundity [10]. Thus, the *ACD6-Est-1 allele* provides both a blessing and a curse, underlying a typical growth-defense trade-off [10]. This allele is nevertheless rather common in natural populations, with about 10% of wild accessions carrying *ACD6*-Est-1-type alleles [9,10], consistent with *ACD6*-Est-1 conferring tangible fitness benefits in nature and balancing selection maintaining the allele despite a growth penalty.

The *ACD6*-Est-1 phenotype shares similarities with the one caused by the EMS-induced gain-of-function mutation *acd6-1*, which also links stunted growth with enhanced immunity [7,8,14,15]. However, *acd6*-1 phenotypes are greatly exaggerated under lower temperatures [9,11], which is typical for many autoimmune genotypes [16], making it unlikely that they will survive outside under the conditions that are typical for much of the natural range of *A. thaliana* [17]. Similar phenotypes as in plants with an *ACD6*-Est-1-type allele are produced by certain heteroallelic combinations at *ACD6*, with the difference that *ACD6* compound heterozygotes, but not *ACD6*-Est-1-dependent necrosis, can be suppressed in the greenhouse by elevated temperature [9,11,12]. Counterintuitively, such heterozygotes can be found in the wild, where temperatures during the typical *A. thaliana* growing season are generally lower than the ones used in laboratory experiments [11]. In addition, when grown outdoors, the fitness of such heterozygotes does not appear to be different from their parental accessions, which has been attributed to microbial load inducing SA and hence reducing growth to a similar extent in the parents as in the hybrid progeny [12].

Here, we sought to further reveal the contribution of hyperactive *ACD6* alleles to plant fitness by characterizing plant phenotypes and microbial associations in both greenhouse and field conditions. Because SA is known to affect the shoot and root microbiota [18–21], we hypothesized that *A. thaliana* plants carrying the *ACD6*-Est-1 allele would have an altered commensal microbial population when exposed to natural microbial sources such as field-collected soil [22–24]. We therefore prepared a set of *A. thaliana* lines with both hyperactive and attenuated ACD6 function, grew the plants in field-collected soil both in greenhouse and field conditions for two consecutive seasons, and monitored molecular, physiological, and microbial phenotypes at flowering. The result was a dramatic display of phenotypic plasticity, with hyperactive *ACD6-Est-1* phenotypes being suppressed in field conditions at both the organismal and molecular level. The prevalence of *ACD6*-Est-1-type alleles in natural populations indicates that they have been maintained by selection, but under which conditions they are beneficial in nature remains elusive. This study not only highlights the risk of studying plant immune system components in controlled conditions only, but it also suggests that a better understanding of the environmental regulation of growth-defense tradeoffs could improve yield in crops that use *ACD6*-like genes as part of their defense repertoire.

## Results

### Phenotypes of *ACD6*-Est-1 plants in the field

To reduce confounding by genetic background without the use of transgenic plants, we took advantage of a heterogeneous inbred family (HIF) derived from an Est-1 x Col-0 cross [25] that was only segregating in the genomic interval around *ACD6* [10]. For simplicity and to minimize bias, we only planted progeny from the heterozygous HIF parents, with the Mendelian expectation of progeny containing approximately 25% individuals homozygous for the *ACD6*-Est-1 allele and 25% homozygous for the standard *ACD6*-Col-0 allele. In addition, we generated transgene-free CRISPR/Cas9 genome-edited Est-1 lines in which *ACD6* was disrupted by a single-base pair deletion [9], resulting in a severely truncated ORF that encodes only the first 37 residues of the 671-amino acid protein. The two contrasts allowed for comparison of *ACD6*-Est-1 and *ACD6*-Col-0 alleles in a mixed, but isogenic Col-0/Est-1 background, and of plants with the *ACD6*-Est-1 allele and an *acd6* knockout mutation (from here on: Est-1:*acd6*-null) in a pure Est-1 background.

We then planted the HIF:Est-1, HIF:Col-0 and Est-1 wild-type genotypes in late fall of 2016 in the greenhouse and at a research site of the University of Tübingen in Tübingen, Southern Germany, in the same batch of topsoil sourced from the Tübingen site. In the greenhouse only, we grew in addition Est-1:*acd6*-null plants in parallel, since genome-edited plants are subject to GMO rules in the EU and therefore cannot be easily grown outdoors without a lengthy permitting process. Instead, we grew Est-1 and Est-1:*acd6*-null plants outdoors in 2019 at a secure facility at Agroscope in Zurich-Reckenholz, Switzerland, using field soil from that site. We started plants from stratified seeds, and the outdoor plants overwintered in an open field without further protection (Methods, Supplementary Figure 1). To harvest faster-maturing greenhouse plants at a similar phenological stage, we started them in late January the following year using remaining homogenized soil that had been left over winter in the field; the greenhouse plants therefore did not pass through a cold vernalization period.

When both indoor and greenhouse plants began to flower at the end of March, we took overhead photos and quantified green pixels of their rosettes (Methods, Supplementary Figure 1) as a proxy for biomass [26]. Greenhouse plants showed typical *ACD6*-Est-1 phenotypes, with HIF:Est-1 and Est-1 being noticeably smaller and more necrotic than HIF:Col-0 and Est-1:*acd6*-null plants (Figure 1a, 1b). Unexpectedly, in the field, late-onset necrosis in HIF:Est-1 and Est-1 was not apparent, and these genotypes were no smaller than their isogenic counterparts with the standard *ACD6*-Col-0 allele or an *acd6* knockout mutation. A repeat planting of the same genotypes in 2017 produced similar results. The following in-depth analyses are mostly based on plants from the first planting season.

**Fig. 1.**
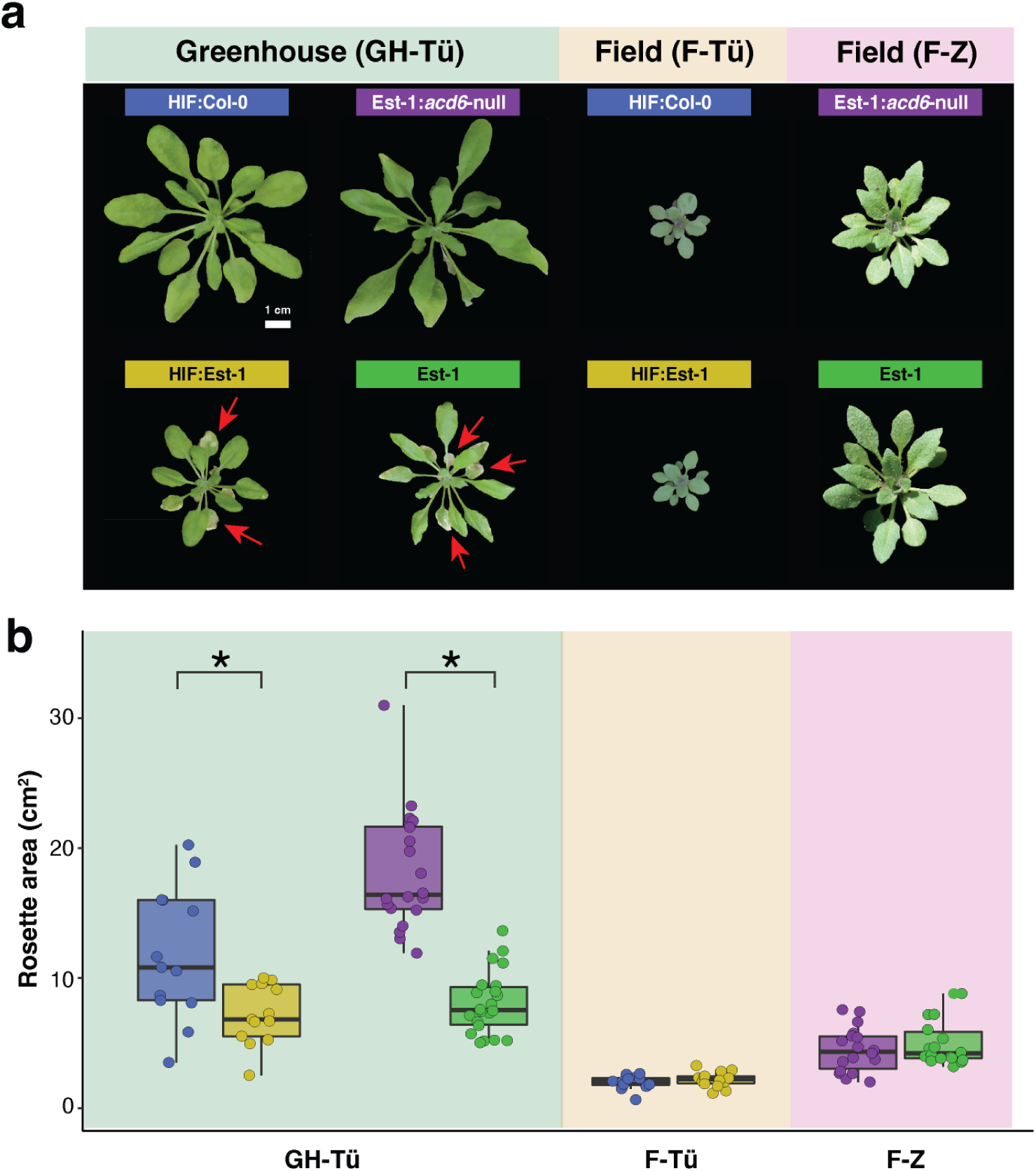
| A hyperactive *ACD6* allele reduces growth and causes necrosis only in the greenhouse. **a**, Representative *A. thaliana* rosettes from different genetic backgrounds. Yellow and blue labels represent descendents of a heterozygous *ACD6* HIF line that are homozygous for either the Est-1 or the Col-0 allele of *ACD6*. Green and purple labels represent the Est-1 accession and the isogenic *acd6* knockout null allele in that accession. Plants were grown in Tübingen in the greenhouse (GH-Tü), outdoors in the field in Tübingen (F-Tü), or outdoors in the field in Zurich (F-Z, a GMO-approved facility). All plants are shown at the same scale (scale bar 1 cm). Note the necrotic older leaves for greenhouse plants with the Est-1 allele (red arrows). **b,** Rosette size of individual plants, calculated by quantifying green pixels from images similar to those in (a). * indicates *P* < 0.01 in a Mann-Whitney U-test.

### *ACD6* drives a strong immunity transcriptional program only in the greenhouse

The lack of visual phenotypes linked to *ACD6* genotype in field conditions suggested that *ACD6* activity is strongly dependent on growth conditions. We therefore extracted mRNA and sequenced transcriptomes from all genotypes in both the field and greenhouse for the 2016 German plants and the 2019 Zurich plants. To avoid transcriptional variance due to circadian factors, we harvested rosettes in a narrow time window, working quickly to minimize wounding responses (Methods). After constructing mRNA-seq libraries with a custom protocol [27], we mapped reads to the Col-0 TAIR10 reference genome (TAIR, http://arabidopsis.org).

We first examined broad-scale differences between the samples through principal component analysis (PCA) ordination, using the 500 genes with the greatest variance across genotypes and environments. Gene expression differed greatly by environment, with greenhouse and field plants cleanly separated (Figure 2a). mRNA levels also differed, as expected, by background genotype, with HIF plants, which have a mixed Col-0/Est-1 background, being distinct from plants in a pure Est-1 background (Figure 2a). In the greenhouse, but not in the field, plants with the hyperactive *ACD6*-Est-1 allele were separated in the PCA from their counterparts with the standard or knockout *ACD6* alleles (Figure 2a). In total, greenhouse conditions led to significant upregulation of 981 genes in HIF:Est-1 as compared to HIF:Col-0, and 1,271 genes in Est-1 as compared to Est-1:*acd6*-null (FDR-adjusted Wald test, *P < 0.01)*. While only 265 upregulated genes overlapped between HIF:Est-1 and Est-1 (Figure 2b), in both backgrounds defense signaling pathways were upregulated, as inferred from Gene Ontology (GO) enrichment analysis (Figure 2c). This is consistent with the previously-observed induction of defense marker genes in plants with *ACD6-*Est-1 and other high-activity *ACD6* alleles [8,10,15]. In stark contrast to the greenhouse, the number of upregulated genes in field plants with the *ACD6*-Est-1 allele was only in the single digits, and they did not overlap between the two genetic backgrounds, and the handful of genes that did differ from the control plants had no known links to the immune response.

**Fig. 2.**
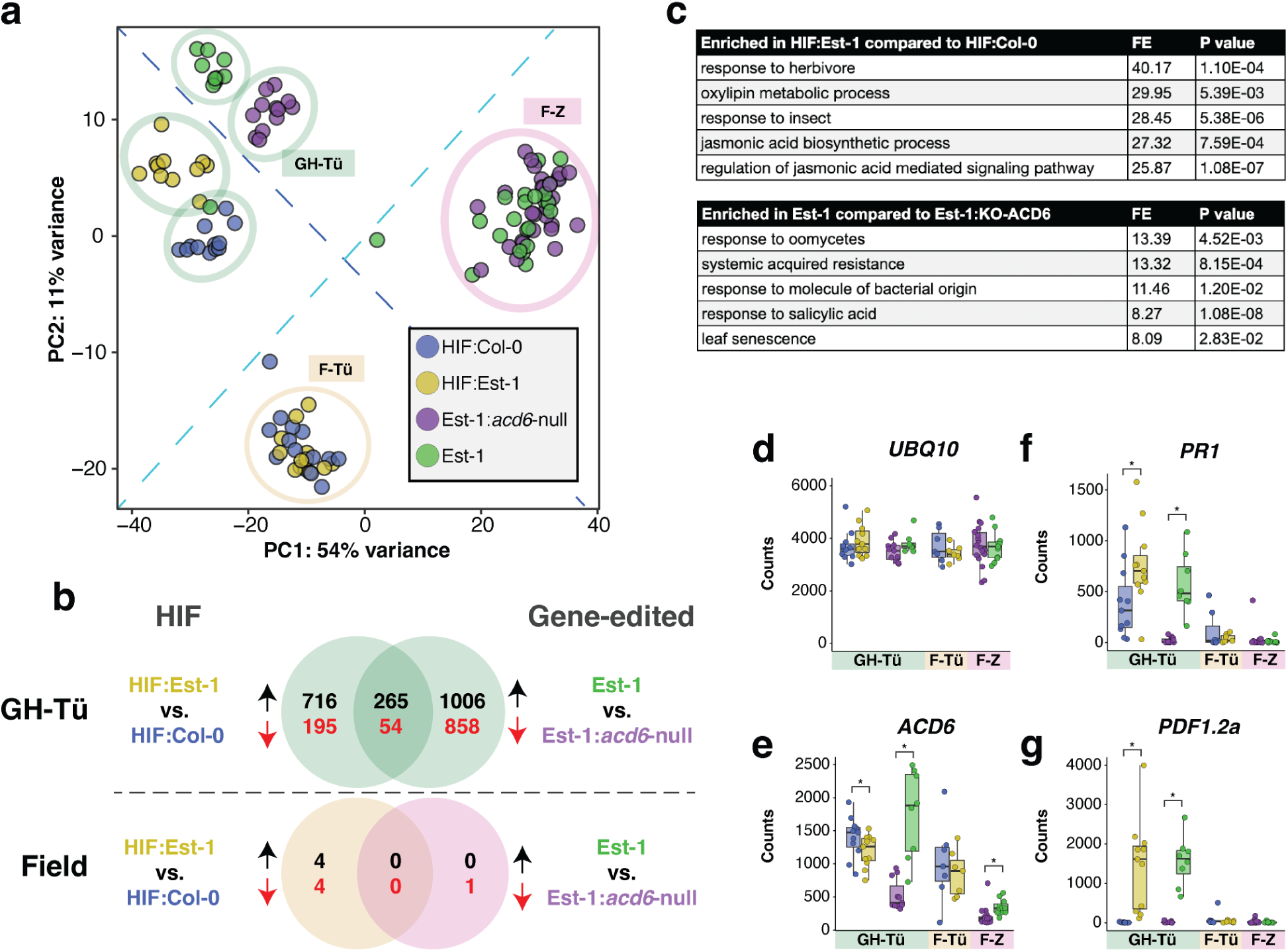
| A hyperactive *ACD6* allele upregulates defense responses specifically indoors. **a**, Principal Component Analysis of mRNA counts calculated from the 500 genes with the most variance across genotypes and environments. Samples are colored by plant genotype, and color code applies also to other panels. Circles indicate the environments (GH-Tü = Greenhouse Tübingen, F-Tü = Field Tübingen, F-Z = Field Zurich), and the color code applies to other panels as well. The dashed light blue line separates field and greenhouse environments, and the dashed dark blue line separates *ACD6* allelic contrasts. **b,** Overlaps between upregulated (black numbers) and downregulated (red numbers) genes in *ACD6* allelic contrasts, either in the greenhouse (top) or field (bottom). **c,** The five gene ontology (GO) processes with the greatest fold enrichment (FE) in *ACD6* allelic contrasts. **d-g,** Normalized mRNA counts for the *UBQ10* reference gene, *ACD6*, the SA marker *PR1*, and the JA marker *PDF2.1a*. Boxes enclose the interquartile range (IQR) with whiskers extending to up to 1.5 times the IQR. * *P* < 0.05 in a Mann-Whitney U-test.

Zooming in on individual genes, we first verified that the commonly-used control *UBQ10* [28] was consistently expressed across our samples (Figure 2d). In both the greenhouse and the field, *ACD6* transcripts in Est-1:*acd6*-null were less abundant than in Est-1, despite the mutation being in the coding sequence, consistent with both nonsense-mediated decay (NMD) as well as positive feedback regulation of *ACD6* [7,29] (Figure 2d). Expression of *ACD6* was also lower in the field compared to the greenhouse, particularly in the pure Est-1 background (Figure 2e). In the HIF background, expression of *ACD6*-Col-0 and *ACD6*-Est-1 alleles was similar for both genotypes, consistent with the results of previous analyses in which either allele had been introduced as a transgene in the Col-0 background [10]. The slightly lower apparent expression of the Est-1 allele in the HIF background could be explained by reduced read mapping of the divergent Est-1 allele to the Col-0 TAIR10 reference.

Classic defense marker genes showed strong differential regulation; the SA-responsive marker gene *PR1* (Figure 2d) and the jasmonic-acid (JA) marker *PDF1.2* (Figure 2e) were greatly upregulated in the greenhouse, but not in the field, in both genetic backgrounds with *ACD6*-Est-1 alleles, consistent with *ACD6*-Est-1 increasing levels of both SA and, more moderately, JA [10]. In the greenhouse, several other marker genes were also more strongly expressed in *ACD6*-Est-1 plants in the greenhouse, but not in the field (Supplementary Figure 2).

We quantified several major plant hormones in greenhouse and field plants by LC-MS. Levels of both SA and its inactive vacuolar storage form SA-2-O-β-D-glucoside (SAG) were higher in greenhouse-grown than in isogenic field-grown plants, and in the greenhouse, more abundant in Est-1 wildtype than in Est-1:*acd6*-null plants, similar to expression of the SA marker gene *PR1* (Supplementary Figure 3, Figure 2). SA and SAG concentrations were not noticeably different in greenhouse-grown HIF:Est-1 plants compared to HIF:Col-0 plants, consistent with the more modest differences in *PR1* expression between these two genotypes in our experiments. Jasmonic acid (JA) accumulated to higher levels in greenhouse-grown than in field-grown plants, and also in greenhouse-grown Est-1 and HIF:Est-1 plants compared to Est-1:*acd6*-null and HIF:Col-0 plants, respectively, mirroring the expression of JA marker gene *PDF1.2a* (Supplementary Figure 3, Figure 2). Indole-3-carboxylic acid (ICA), a tryptophan-derived indolic hormone implicated in defense [30], showed a similar pattern to JA with increased levels in greenhouse-grown plants carrying *ACD6*-Est-1 and relatively lower accumulation in field-grown plants. Indole-3-acetic acid (IAA), the only hormone tested with merely peripheral roles in defense, was also the only hormone with higher levels in field-grown plants, and the only that appeared unaffected by *ACD6* genotype.

### The Est-1 allele of *ACD6* does not create an obvious fitness liability in the field

The mRNA, plant hormone phenotypes, and striking visually-apparent *ACD6*-dependent differences in both plant size and late-onset necrosis strongly suggested that there was a sustained higher level of *ACD6* activity in the greenhouse not only at harvest but also throughout the plant life cycle, which should affect plant fitness. The lack of clear *ACD6*-dependent phenotypes in the field plants, however, did not exclude the possibility that *ACD6* was nonetheless affecting field plant fitness, e.g., by affecting seedling survival or pollination success. This concern was especially relevant because our harvest timepoint at the onset of flowering was not suited to address any role *ACD6* might play during later stages of the life cycle, including flowering and seed set.

To determine the contribution of *ACD6* alleles to reproductive fitness, we collected first generation seeds (G1) from HIF parents heterozygous for the *ACD6*-Col-0 and ACD6-Est-1 alleles in three starting batches. As a control, we first germinated seed aliquots of each in bulk on agar, extracted DNA from pooled seedlings and determined allele ratios by amplicon sequencing; this analysis confirmed that the germinated G1 seedlings had equal representation of *ACD6* alleles (Methods, Figure 3b). We next planted hundreds of G1 seeds at high density in plastic trays and germinated them in the greenhouse and also in two outdoor field environments around Tübingen, allowing all survivors to grow to maturity. After collecting second generation (G2) seeds in bulk, we determined the *ACD6* allele ratio in the viable seeds by germinating G2 seeds on agar and measuring the *ACD6* allele ratios (Figure 3b). In the field, the ratio of *ACD6*:Est-1 to *ACD6*:Col-0 alleles remained balanced. However in the greenhouse, the *ACD6*:Col-0 allele was twice as common as the *ACD6*:Est-1 allele, demonstrating a clear and strong fitness penalty for the Est-1 allele in the greenhouse (Figure 3c), consistent with the considerably smaller stature in that environment (Figures 1 and 2).

**Fig. 3.**
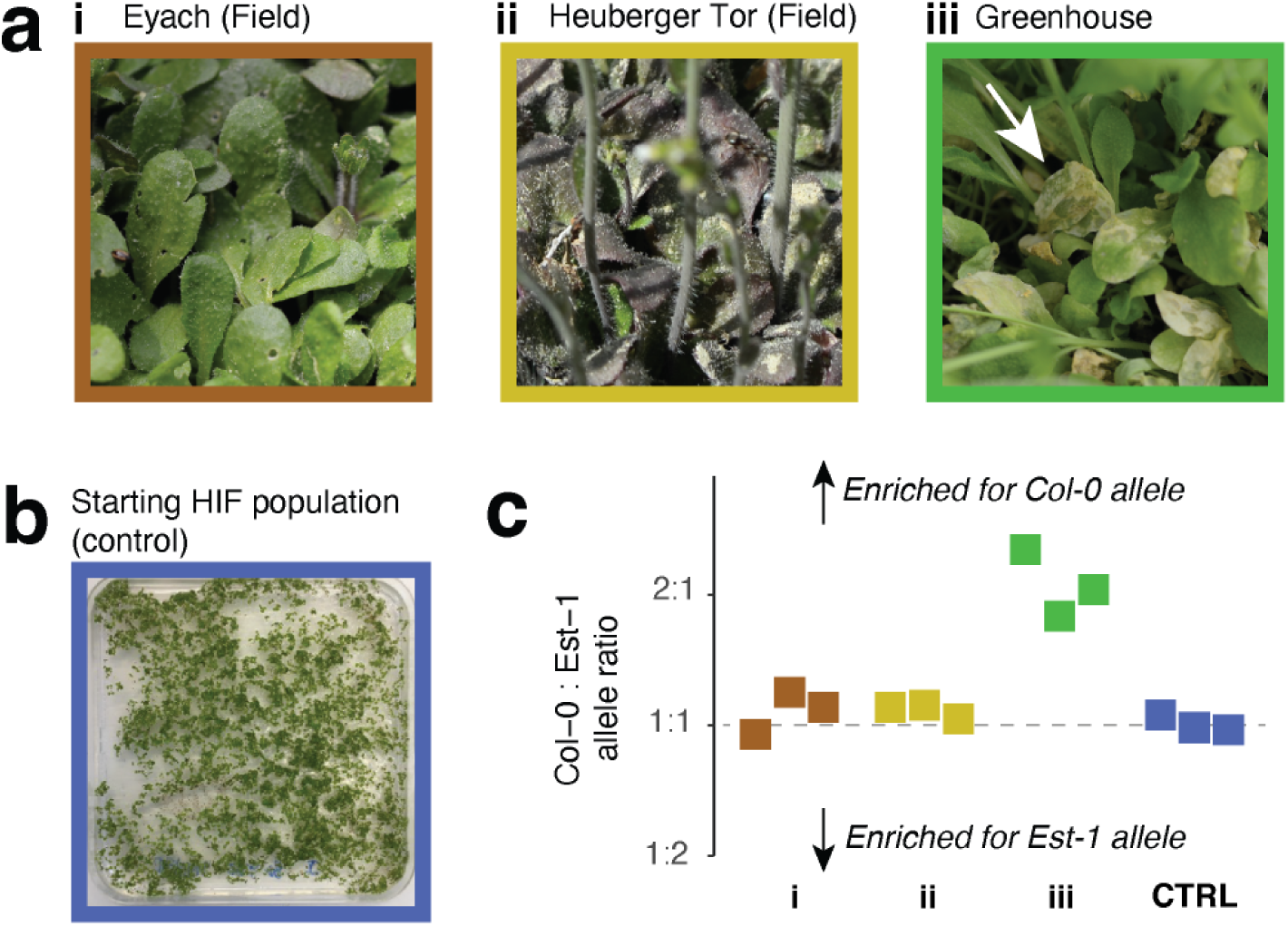
| The fitness penalty of *ACD6*-Est-1 is limited to the greenhouse. **a**, Leaves of HIF progeny raised to maturity at high density in different environments. **i:** Flowering plants in the field at Eyach, with some leaf holes due to insect herbivory. **ii:** Flowering plants in the field at Heuberger Tor, with darker leaf colors due to anthocyanin production. **iii:** Flowering plants raised in the greenhouse. Note the necrotic leaves (arrow) characteristic of plants carrying *ACD6*-Est-1. **b,** Seedlings grown from the same batch of seeds used in (a), germinated on agar as a control to capture the allele frequency in the starting population. **c,** Ratio of *ACD6*-Col-0 to *ACD6*-Est-1 alleles recovered in bulk from freshly-germinated viable seeds collected from mature plants from (a) or from the starting population in (b). Colors represent different environments as indicated by the picture border colors in panels (a) and (b). The vertical dashed line represents a balanced allele ratio of 1:1.

### Temperature does not explain difference between greenhouse and field conditions

A sustained greenhouse temperature of 23°C is higher than the average temperature *A. thaliana* usually experiences in the field during its main growth period [17,31]. Further, many ecologically relevant phenotypes in *A. thaliana*, including the activity of immune system, are often affected by shifts between 23°C and 16°C [11,32,33]. We therefore considered that the sustained higher temperatures in the greenhouse might induce the deleterious effects of *ACD6*-Est-1, even though lower temperature normally enhances rather than ameliorates immunity-dependent phenotypes [16]. However, in controlled conditions, at both 23°C and 16°C, we observed clear developmental differences between Est-1 and Est-1:*acd6*-null plants (Supplementary Figure 4). We conclude that environmental factors other than simply a change in temperature are responsible for the phenotypic differences in expression of the *ACD6*-Est-1 phenotype between greenhouse and the field.

### *ACD6*-Est-1 does not affect the foliar microbiome in the field

While our molecular, metabolic and phenotypic analyses did not reveal any obvious differences between mature plants with and without the *ACD6*-Est-1 allele in the field, we hypothesized that this could have been due to *ACD6*-Est-1 only acting in early seedlings or intermittently in field conditions. If this were the case, there might be a footprint in the microbiota that colonizes plants with an *ACD6*-Est-1 allele.

For the analysis of the bacterial microbiota, we prepared and sequenced bacterial amplicons of the V4 region of the 16S rRNA region (hereafter rDNA) from total leaf DNA extracted from rosettes of field– and greenhouse-grown rosettes n Tübingen in 2016-2017, a biological replicate in Tübingen 2017-2018, and a single field replicate of the Est-1 vs. gene-edited Est-1:*acd6*-null lines grown outdoors in Zurich in 2019. We compared the results with previously-generated V4 16S rDNA data from the phyllospheres of wild *A. thaliana* populations collected near Eyach in the Tübingen region during a previous season [34]. For fungi, we generated internal transcribed spacer (ITS) data from rosettes collected in Tübingen in 2016-2017, comparing these previously-generated ITS data from the natural Eyach population [34].

We first classified amplicon sequence variants (ASVs) from all datasets to the family level and clustered samples by similarity. Unexpectedly, the bacterial microbiomes of the plants from the German field experiment were much more similar to that of wild *A. thaliana* plants with different genetic backgrounds, growing in diverse soils, being of a wider range of rosette size and developmental status and harvested in a different year than than they were to the microbiomes of isogenic greenhouse plants grown in the same soil and harvested at the same time (Figure 4a). This was also the case for fungal microbiomes (Figure 4b), demonstrating the smaller role of soil compared to other environmental factors in structuring outdoor foliar microbiomes. When we looked specifically in field plants for ASVs that differed in abundance between HIF:Est-1 and HIF:Col-0 plants, or in Est-1 and Est-1:*acd6*-null plants, we found none. Even in the greenhouse, using the very same plant material for which we had already demonstrated substantial differences in expression of immunity markers due to *ACD6* alleles (Figure 2), we failed to find differentially-abundant bacterial or fungal ASVs that distinguished plants with or without *ACD6*-Est-1. This was particularly surprising because SA, the levels of which are highly sensitive to *ACD6* activity [11,12,14], has been shown to alter the microbiota in many other settings – although these effects are clearer in roots than in leaves [18–21].

**Fig. 4.**
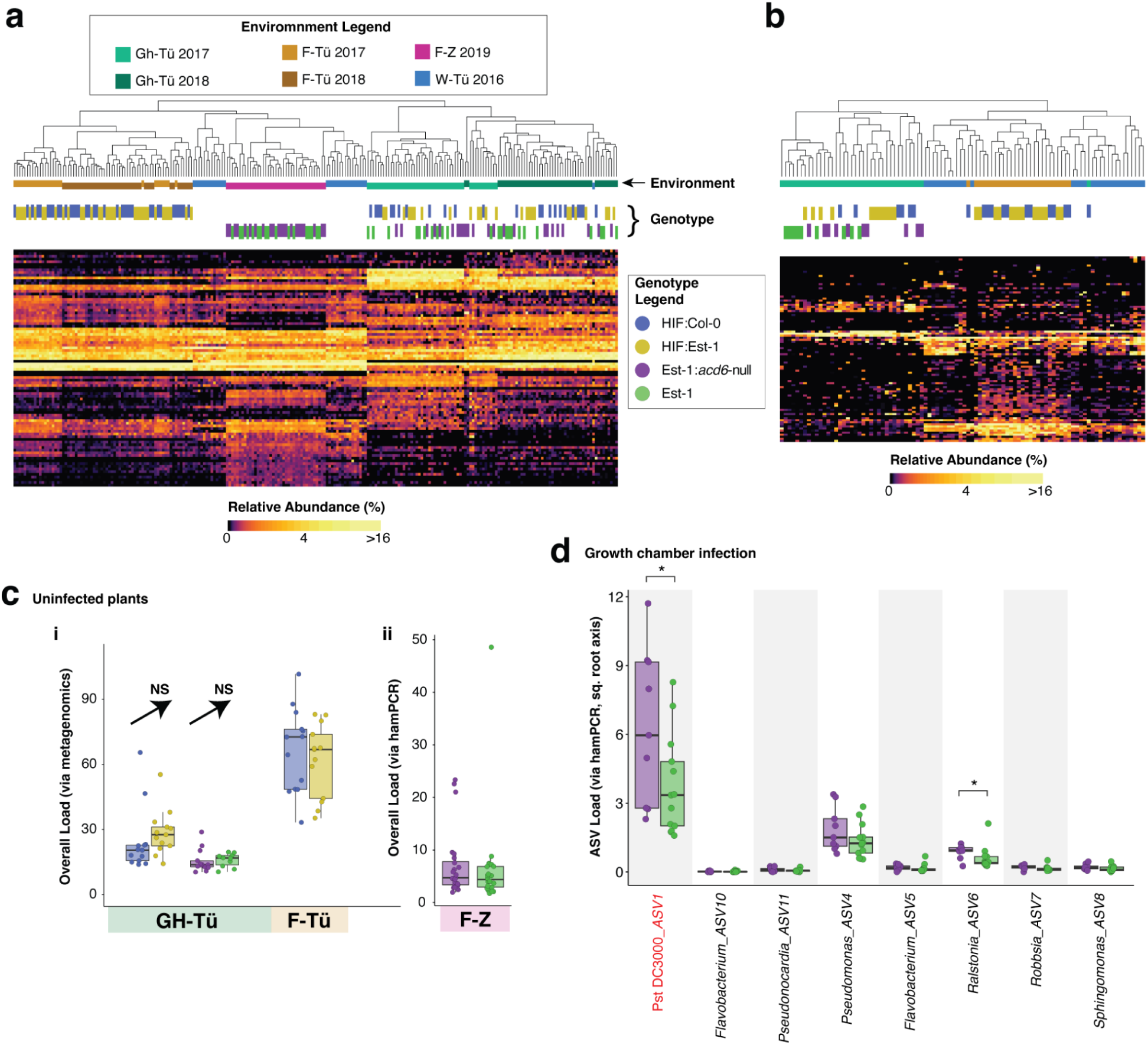
| *ACD6*Est-1 has negligible effects on colonization by a natural microbiome. **a**, Heatmap showing square root transformed relative abundances of bacterial families (rows, unlabeled) based on classification of V4 16S rDNA bacterial reads from both planted *A. thaliana* individuals and plants samples from wild populations (columns in heatmap). Samples in the heatmap are clustered by Bray-Curtis dissimilarity, with groupings shown in the dendrogram. The environment of cultivation is indicated by a horizontal color key (vertically staggered only to improve visibility) between the dendrogram and heatmap, using colors in the “environment legend”. The genotype of the plants is similarly shown using colors in the “genotype legend”. For the environment legend, Gh-Tü = greenhouse Tübingen; F-Tü = field Tübingen; F-Z = field Zurich; W-Tü = wild Tübingen (natural populations near Tübingen). **b,** Same as (a), but showing fungal families based on ITS2 amplicons. **c,** Bacterial load, **i:** for Tübingen-grown plants from 2017 as calculated by the ratio of bacterial reads to plant reads in metagenome data. **ii:** for Zurich--grown plants from 2019 as deduced from hamPCR. Environment and genotype colors follow the same code as in (a). Boxes enclose the interquartile range (IQR) with whiskers extending to up to 1.5 times the IQR. **d,** Bacterial loads of the most abundant ASVs in growth chamber-grown plants infected with Pst DC3000 and a mix of uncharacterized phyllosphere microbes. \**P* < 0.05 in a Mann-Whitney U-test. The ASV corresponding to Pst DC3000 is labeled in red.

We next reasoned that perhaps the broad-scale immune response triggered by *ACD6* might reduce the colonization of all microbes rather than select taxa, and we therefore compared the ratio of microbial sequencing reads to plant reads in whole-genome shotgun data [34] or hamPCR data [35] to calculate microbial load in our samples (Methods). While this revealed field plants to have approximately double the bacterial load as greenhouse plants, there were no *ACD6*-dependent differences in microbial load either in the greenhouse or the field (Figure 4c).

Given these negative results, we reasoned that *ACD6*-dependent defenses may be able to restrict specifically the growth of invasive opportunistic microbes without affecting the commensal microbes that colonize undamaged surfaces. We therefore conducted *Pseudomonas* infections in both the field and greenhouse. For the field infection, we grew overwintered HIF plants outdoors in Tübingen, sprayed them in the spring prior to bolting with a mixture of four Pv-ATUE5 strains that occur naturally on *A. thaliana* in this geographic region [36,37], and harvested plants one week later. As in all other experiments with field-grown plants, we noticed no *ACD6*-dependent differences in size or in late-onset necrosis. This time, unlike prior harvests, we then surface-sterilized all rosettes as in [38] to remove any transient or loosely-unassociated microbes, and sequenced 16S rDNA amplicons using hamPCR to also determine bacterial load [35]. Again, we found no differences in bacterial associations with HIF:Est-1 vs. HIF:Col-0 homozygotes, including for the ASV corresponding to Pv-ATUE5 (Supplementary Figure 5a), and we further observed no *ACD6*-dependent difference in bacterial load (Supplementary Figure 5b). For the greenhouse infection, we grew Est-1 and Est1:*acd6*-null plants for five weeks in short days in a growth chamber and sprayed them with either *Pseudomonas syringae* pv. tomato DC3000 (Pst DC3000), an uncharacterized mixture of cultured *A. thaliana* phyllosphere microbes (“other” microbes), both Pst DC3000 and “other” microbes, or vehicle (MgCl_2_). We harvested 3 days post infection and determined bacterial load and composition with hamPCR. Here, while there was no difference in total microbial abundance caused by *ACD6* genotype (Supplementary Figure 5c), we did observe genotype-dependent differences in abundance for a handful of the more abundant ASVs (Figure 4d). Importantly, this included the ASV corresponding to Pst DC3000, which had a significantly higher load in Est-1:*acd6*-null than in Est-1 plants (Figure 4d, Supplementary Figure 5e), consistent with reduced immune function when ACD6 function is lost completely (Lu et al. 2003; Zhu et al. 2021; Tateda et al. 2014). For plants sprayed only with Pst DC3000, we also enumerated Pst DC3000 colony forming units (CFUs); this orthogonal approach also revealed, in support of the hamPCR data, a slightly higher pathogen load in Est-1:*acd6*-null plants (Supplementary Figure 5d). Besides Pst DC3000, an ASV corresponding to a Ralstonia strain was also significantly more abundant in Est-1:*acd6*-null plants.

## Discussion

The *ACD6* gene was originally characterized through a lab-induced gain-of-function *acd6*-1 mutation that increases levels of SA, spontaneous cell death, and resistance to pathogens, but also reduces plant stature [14]. *acd6*-1 is an example of lesion mimic mutants, which exhibit disease-like symptoms in the absence of pathogens [39]. Such mutants have been described in many species, especially in maize, where they represent the largest class of gain-of-function mutants [39]. The lesion mimics phenotype is often strongly environmentally-dependent, and *acd6-1* is no exception, with the phenotype much more severe at 16°C and attenuated at 23°C [9,11].

In general, while induced lesion mimics not only provide insight into gene function, plant immunity, and plant physiology, they are also useful to understand trade-offs between growth and disease resistance. However, it is unclear whether natural lesion mimics might confer long-term evolutionary benefits. For example, the *acd6*-1 mutant did not survive in our field conditions, and other mutants, such as *cpr5*, that constitutively increase SA levels, greatly reduce fitness in field conditions [40]. What is fascinating about the *ACD6* locus is that natural alleles, such as Est-1 type alleles, have a hyperactive immune system that is in many ways similar to the lesion mimic mutant *acd6*-1, yet they do not have the same temperature sensitivity (Supplementary Figure 4) [9,11]. That *ACD6*-Est-1 alleles, which differ substantially in sequence from standard *ACD6* alleles, are maintained in wild populations at substantial frequencies, about 10-20%, [10,11,13] is indicative of *ACD6* allelic variation being under balancing selection in nature – a hallmark of many though not all immune genes [41–46]. Given the documented roles of *ACD6* in immunity under greenhouse conditions, the most parsimonious explanation is that balancing selection at *ACD6* is best explained by increased defense outweighing the drawbacks of reduced growth. We envisaged a simple scenario, wherein a natural hyperactive *ACD6* allele would result in increased SA levels in field plants, just as it does in greenhouse plants, and that this would lead to obvious phenotypic consequences in the field. However, the reality appears much more complex.

In our field experiments, we could neither observe a benefit nor a cost of the *ACD6*-Est-1 allele, and we observed no major differences between HIF lines that differed in *ACD6* genotype in either life-time fitness or markers of immune system activation. Although we were not able to measure field fitness in Est1:*acd6*-null plants, which would require letting plants set seeds, the fact that we did not observe any other phenotypic difference between Est-1 and Est1:*acd6*-null plants in the field in Zurich suggests that life-time fitness was also unaffected even when compared to a complete loss of *ACD6* activity. Further supporting our observations, it was previously shown that heterozygous combinations of certain *ACD6* alleles in the same plant, including combinations that naturally occur in hybrid zones, can lead to severe hybrid necrosis in the greenhouse, but that these are partially attenuated in field conditions [11,12]. Therefore, we speculate that *ACD6* hyperactivity in nature is very context dependent, being expressed either only during very specific developmental stages or only under very specific environmental conditions.

We do not know yet what aspect of the greenhouse environment results in such strongly variable expression of *ACD6* hyperactivity. One obvious possibility is that the controlled greenhouse has less abiotic stress such as drought as well as fluctuating insolation and temperature; abiotic stress is known to suppress plant immune responses [47]. Indeed, that the greenhouse had more favorable conditions can be inferred from more plants growing more quickly and much larger (Figure 1). The most upregulated mRNA GO terms in the field compared to the greenhouse were “response to light intensity” and “carotenoid synthesis”, so it is plausible that light stress in the field was a factor in suppressing SA pathways [48]. Another possibility is that greater outdoor UV-B radiation prompted greater production of protective secondary metabolites, reducing the need for SA-mediated pathways [48]. A further possibility is that *ACD6* hyperactivity is growth rate dependent, with the slower growth outdoors greatly dampening *ACD6* hyperactivity. In this scenario, *ACD6* only comes into play under circumstances where plants encounter very favorable circumstances for growth. Yet another possibility is that one or more members of natural microbial populations on the field plants suppress SA-mediated defenses, including *ACD6* [49].

It is important to note that in natural accessions, the activity of *ACD6* alleles also depends on variation at other loci [10,13]. The small proteins MHA1 and MHA1-like (MHA1L) can interact with ACD6 to suppress or enhance its activity [9]. The full details of how such cryptic modifiers of *ACD6* activity respond to fluctuating environmental cues are unknown, and these loci could be important in the phenotypic plasticity observed here.

We also tested the hypothesis that transient hyperactivity of *ACD6-Est-1* in the field might have lasting effects on the foliar microbiome. Using amplicon sequencing, we did not observe differential natural microbial colonization across *ACD6* genotypes, even in greenhouse conditions where *ACD6*-dependent phenotypes were very clear and conditions for which an A*CD6*-dependent reduction in accumulation of pathogenic microbes has been previously documented [10]. Using both shotgun sequencing and hamPCR, we also did not observe an effect of *ACD6* on bacterial load, even in the greenhouse. We addressed this confusing situation by inoculating greenhouse plants with commensal microbes in the presence of a bacterial pathogen and looking at both pathogen and commensal abundance via amplicon sequencing. Both colony counting and hamPCR revealed a slight but statistically-significant reduction in the proliferation of Pst DC3000 in Est-1 compared to Est1:*acd6*-null plants, consistent with previous observations [7,9,50]. However, few other ASVs seemed to be affected. Our data strongly suggest that, despite very high SA levels, effects of the *ACD6*-Est-1 allele on commensal phyllosphere bacterial and fungal microbes are slight. To date, few studies have addressed the role of SA levels on foliar microbes, and the clearest effects have been observed in hyperimmune mutants with extreme phenotypes grown in gnotobiotic conditions with a synthetic community of bacteria [20,51]. We propose that the suite of broad-spectrum defenses upregulated by *ACD6* behave like a “speed bump” in traffic, which poses little hindrance to vehicles maintaining the speed limit, but which seriously hinders or damages vehicles traveling too fast. Similarly, commensal microbes that grow slowly without damaging the plant may be able to tolerate increased intracellular concentrations of antimicrobial peptides and secondary metabolites that they rarely encounter, but pathogens that rely on extraction of plant resources for fast growth are much more sensitive to such defenses.

The negative impact of the natural *ACD6*-Est-1 allele on seed number of greenhouse-grown plants was severe. Should such *ACD6* alleles be activated at the wrong times in field conditions, even slightly, this could have significant effects on fecundity. Knowing the triggering environmental conditions, how they modulate *ACD6*-Est-1, and how important different *ACD6* alleles are for fitness during epidemics could have practical relevance for major cruciferous oilseed crops like *Brassica napus*, for which *ACD6*-orthologs can be identified bioinformatically, and for which there is a wide bank of potential breeding germplasm.

## Methods

### Plant material and seed treatment before planting

*Arabidopsis thaliana* accessions and heterogeneous inbred family (HIF) lines were derived from stocks maintained in the lab. All seeds were surface-sterilized by a 1 mine submersion in 70% ethanol with 0.01% Triton X-100, followed by a 12 min submersion in over-the-counter bleach (DanKlorix; 2.8% NaOCl w/w) diluted to 10% in water, followed by three washes in sterile distilled water. Seeds were left in water and stratified at 4°C in sterile centrifuge tubes for 1 week before planting.

For genome editing, an *A. thaliana* codon-optimized Cas9 (*athCas9*) [52] was used, with the final *pUBQ10::athCas9:trbcs::gRNA::mCherry* constructs assembled from six GreenGate modules [53]. The same gRNA was encoded on the transgene by 5’-GTGTC GCCCG TAGGT GACG-3’for both ACD6-Col-0 and ACD6-Est-1. The primers for generating the gRNA cassette were 5’-GTGTC GCCCG TAGGT GACGG TTTTA GAGCT ATGCT GAAA-3’ and 5’-CGTCA CCTAC GGGCG ACACC AATCA CTACT TCGAC TCTA-3’. Red fluorescence from a *pAT2S3::mCherry:tMAS* cassette [54] was used for selection of transgene-free seeds. The edited lines were whole-genome-sequenced using 2×150 bp reads on the HiSeq3000 platform (Illumina).

### Field cultivation in Tübingen

Topsoil was obtained from a research field of the University of Tübingen at the Heuberger-Tor-Weg in Tübingen, Germany (48°32’44.9”N 9°02’32.8”E) in October 2016 for most plants, and again in 2017 for a replicate experiment. The soil was initially spread out in plastic trays in a plastic foil tunnel and allowed to dry for one week until the clayey soil began to crumble to the touch, which made it easier to break larger aggregates in the soil with physical force and to sieve the material to make it more homogeneous. All soil was sieved through a 8 mm mesh to remove rocks and twigs, mixed thoroughly by repeatedly turning the pile over with a shovel (Supplementary Figure 1), and filled evenly into 40-pot quickpot “QP 40 T/11.5” plastic trays (HerkuPlast Kubern GmbH, Ering, Germany) that were placed inside flat trays to catch water from the top, either rain outdoors or from top watering. After pots were filled, the remaining soil was transferred into 10 L plastic pots for storage and overwintering outside until it could be homogenized again for use in the later greenhouse experiment (see below: “Greenhouse cultivation in Tübingen”).

We pre-watered each 40-pot flat with distilled water by bottom watering and sowed surface-sterilized, cold-stratified seeds at the end of October 2016 (and again in October 2017) when daily highs were around 14°C and lows around 1°C. Stratification for 1 week prior to sowing ensured uniform germination of the cohort, coinciding with the germination of wild winter-annual *A. thaliana* plants in local stands around Tübingen. Each plant of the HIF genetic background was planted in its own pot. Since individual seeds had not been genotyped, plants were automatically randomized for *ACD6* genotype. After germination outdoors for 1 week under transparent plastic lids, the flats spent another month without lids in an open foil tunnel [55] allowing exposure to wind and ambient temperatures but protection from precipitation. Wild plants of other species that germinated, as well as excess *A. thaliana* seedlings, were removed regularly with tweezers so that a single seedling developed in the middle of the pot. Finally, after the first true leaves appeared, plants were moved to an open location where they had full environmental exposure, and the drip trays were removed to prevent water accumulation and flooding during rainfall.

### Field cultivation in Zurich

For CRISPR/Cas9 mutant plants, we followed a similar protocol as above to stratify and germinate Est-1 and Est-1:*acd6*-null seeds at the end of October 2018 in local Swiss soil at Agroscope in Zurich-Reckenholz, Switzerland (47°25’40.8”N 8°31’01.1”E), in a secure open-air room. Instead of placing pots in flats, they were arranged on open soil growth boxes, and extra soil was filled around the pots to hold them in place. Overwintered seedlings were cleaned of weeds and thinned to 1 per pot on 21 February 2019 and finally, mature plants were harvested on 21 March 2019.

### Greenhouse cultivation

Field soil used for greenhouse cultivation was prepared as described above beginning in autumn, with the portion used for field cultivation distributed into pots immediately and the portion to be used for greenhouse cultivation placed in 10 L open plastic pots for storage and overwintering. In January 2017, these 10 L pots were emptied into a single pile which was re-homogenized by turning it over repeatedly with a shovel, and the soil was then filled into 40-pot quickpot plastic trays that were in turn placed inside flat trays to catch water, exactly as for field cultivation. On 1 Feb, 2017 these trays were placed in a greenhouse room without any supplemental lighting averaging 24°C daily highs and 20°C nightly lows, and seeds were planted as above. After 1 week of germination under plastic lids, the lids were removed and the greenhouse plants were top watered with distilled water from a watering can to mimic rainfall including water splash from the soil onto the plants. Mature plants were harvested on 22 March, 2017.

### Growth chamber cultivation

After stratification, seeds were germinated and cultivated in growth rooms at a constant temperature of 23°C or 16°C, air humidity at 65%, 16 hr (long days) day length. Philips GreenPower TLED modules (Philips Lighting GmbH, Hamburg, Germany) provided 110 to 140 µmol m^-2^ s^-1^ light with a mixture of 2:1 DR/W LB (deep red/white mixture with ca. 15% blue) and W HB (white with ca. 25% blue) modules, respectively.

### Imaging and harvesting plants

Both greenhouse and field plants in all locations began to develop a floral meristem at the end of March, and thus were phenologically similar at the time of harvest. We first took overhead photos of all plants using a LUMIX DMC-TZ71 digital camera (Panasonic, Osaka, Japan) without flash. Because of slight differences in the focal length between photos, all images were rescaled in Adobe Photoshop against an internal standard such that each pixel represented the same true area. For smaller field plants where the rosettes were fully contained within the circumference of their pot, a predefined mask was used to extract the area surrounding each plant in the image, and automatic segmentation based on pixel color was applied to distinguish plant leaves from background. For larger greenhouse plants, where leaves had projected beyond the perimeter of the pot, we extracted the area surrounding each plant and manually erased leaf tips from neighboring plants in Adobe® Photoshop® before using automatic segmentation to recognize green pixels of the central plant in each image. Rosette area was then approximated as green pixels in each image.

In all locations, plants were harvested in a randomized fashion between 10:30 am and 12:30 pm to minimize circadian effects. In Tübingen, field plants were harvested first, and greenhouse plants on the following day. All rosettes were first removed from their roots with scissors and tweezers that had been dipped in 95% ethanol and flame-sterilized, and rosettes were placed into 50 mL tubes for washing. Soil and dried mud (potentially containing irrelevant unassociated soil microbes) were washed off leaf surfaces by adding approximately 25 mL of non-sterile distilled water, shaking by hand, and then decanting. This was repeated until the decanted water was visually clear. Next, the rosette was washed once with sterile (autoclaved) distilled water using the same procedure. Finally, the rosette was removed from the tube with re-sterilized tweezers and placed on autoclaved paper towel for blot drying. The clean and dry rosette was loaded into a 2mL or 15 mL screw-cap tube (Sarstedt, Nümbrecht, Germany) depending on the plant size, and snap-frozen in liquid nitrogen. The entire harvest process from cutting to freezing took place in under 3 minutes for each rosette to minimize transcriptional noise from harvest-related stress.

### Homogenization of plant material for DNA, RNA, and phytohormone extraction

For rosettes in 15 mL tubes, we first added 5 autoclaved 5 mm hardened steel balls (VWR, Radnor, USA), re-froze the tubes with the rosettes, and shook the deep-frozen tubes by hand to break the rosettes down into particles of about 1 mm. We then held the steel balls to the cap using an external magnet, and tapped approximately 0.25 mL of the frozen plant powder into an open and pre-frozen 2 mL tube. To these 2 mL tubes, and also to rosettes already in 2 mL tubes, we added 0.5 mL of 1 mm garnet rocks (BioSpec Products Inc., Bartlesville, USA) and a 5 mm glass ball (Carl Roth, Karlsruhe, Germany) and immediately re-froze them to prepare for bead beating. We crushed the frozen plant tissue “dry” in a FastPrep 24 5G homogeniser (MP Biomedicals, Eschwege, Germany) at 4 m/s for 20 s, and immediately re-froze the resulting powder.

### DNA and RNA co-extraction

To approximately 250 mg of frozen plant powder, prepared as described above, we added 750 µL of RNA/DNA lysis buffer (100 mM Tris pH 8.0, 100 mM NaCl, 10 mM EDTA, 1.5% SDS, 2% 2-mercaptoethanol, and 100 µg/mL Proteinase K) that had been pre-warmed to 50°C to enable the buffer to reach all frozen plant particles and to inactivate RNAses. We then did a final high-speed “wet” bead beating at 6 m/s for 60 s to lyse all cells. The tubes were centrifuged at 10,000 x g for 5 min. Of the approximately 600 μL supernatant, 450 μL was transferred to a 1.5 mL centrifuge tube for DNA prep, while 150 μL was transferred to another 1.5 mL centrifuge tube for RNA extraction. For DNA extraction, the 450 μL lysate was mixed with 150 μL sterile 5 M potassium acetate in a 1.5 mL centrifuge tubes to precipitate the SDS. The tubes were spun at 10,000 x g for 5 min and the supernatant transferred to a new 1.5 mL tube. The resulting supernatant was centrifuged a second time to clear out remaining plant material and precipitate, and 600 μL was transferred to a 1 mL deepwell plate (Mettler Toledo, Gießen, Germany). Finally, 360 μL SPRI beads were added to 600 μL of the supernatant (0.6: 1 ratio). After mixing and incubating on a 96-well Magnet Type A (Qiagen, Hilden, Germany), the beads were cleaned with 80% ethanol and DNA was eluted in 100 μL EB (10 mM TRIS, pH 8.0).

### RNA-seq

Between 500 ng and 1,000 ng total RNA was used to construct libraries as described in [27]. Briefly, we used oligo dT) beads to purify plant mRNA, performed first– and second-strand cDNA synthesis, added adapters by ligation, and amplified the library molecules with 12 cycles of PCR before sequencing with 150 bp short reads on an Illumina HiSeq 3000 instrument. We mapped all transcripts to the Col-0 reference genome (TAIR 10) using RSEM [56] and analyzed count tables with the “DeSeq2” package in R [57]. Genes were considered to be differentially expressed if they had an FDR-adjusted *P* value less than 0.01 in a Wald test.

### Phytohormone measurements

Between 50 mg and 150 mg of frozen plant powder was shipped on dry ice to the Mass Spectrometry Core laboratory at the University of North Carolina. LC-MS quantification was performed using a PE Sciex 3000 mass spectrometer equipped with a CTC autosampler and Shimadzu LC system.

### Genotyping of *ACD6*:Est-1 and *ACD6:Col-0* alleles in HIF lines

#### High throughput screening of allele ratios in pooled HIF seeds

Seeds were surface sterilized for 1 min with 70% EtOH and 0.01% Triton X-100, followed by 12 min in 10% bleach, and finally three washes with sterile water. They were stratified for 1 week at 4°C, and germinated on 1% agar containing half-strength MS medium and 5 mM MES buffer. Seedlings were germinated under 16 hours light at 23°C and 65% relative humidity under cool white fluorescent light of 125 to 175 μmol m^-2^ s^-1^ and grown for 1 week, allowing cotyledons to emerge. Liquid nitrogen was then poured on the plates to snap freeze all plant material, and frozen, brittle cotyledons were raked off by spatula and pooled. The pooled plant material was further homogenized with a mortar and pestle, and an aliquot of approximately 250 mg macerate was used for DNA extraction as described for DNA in “DNA and RNA co-extraction”.

The following primers were used to amplify a region of ACD6 containing polymorphisms distinguishing ACD6:Est-1 from ACD6:Col-0, and sequenced using an Illumina MiSeq 50 bp kit. 5’-<u>ACACT CTTTC CCTAC ACGAC GCTCT TCCGA TCT</u> tgtac tctta tttgg gcgca gtt-3’ (>Genotype_ACD6_F) and 5’-<u>GTGAC TGGAG TTCAG ACGTG TGCTC TTCCG ATCT</u> aactc agata tgtct atagt cagca ta-3’ (>Genotype_ACD6_R). In these primers, the lowercase region is complementary to the ACD6 template, while the uppercase region adds overhangs to which Illumina Nextera primers can bind in a subsequent PCR round. First-round PCR with the ACD6-complementary primers used the following program: 95°C for 2 min, 30 cycles of 95°C for 15 s, 60°C for 20 s, and 72°C for 30 s, followed by a final 5 min at 72°C. Illumina adapters were added as described in [34] for “Batch 3 plants”. The full amplicon, including Illumina adapters, was 366 bp long and the sequenceable region was 299 bp. 50 bp of sequencing from the forward primer was sufficient to confidently genotype the Col-0 vs. Est-1 alleles as shown below, where lowercase letters represent the bases corresponding to the primers above and capital letters represent the amplified region:

ACD6_Col-0: tgtac tctta tttgg gcgca gtt **G**GGTG ATC**T**A **GCA**CT CATCC **T**CAAA TC

ACD6_Est-1: tgtac tctta tttgg gcgca gtt **A**GGTG ATC**C**A **AAC**CT CATCC **G**CAAA TC

#### Determining genotypes of individual HIF plants at ACD6

For individual HIF plants, high-quality DNA was prepared. Two reverse primers, 5’-CAAAA CAAGT TTTGA TCTTA CG-3’ (acdHIF_estR G-47730) and 5’-GCCGC TTCTC AGAGC TAG-3’ (acdHIF_col R G-47731), diagnostic for Col-0 and Est-1 sequences were used individually in combination with the same forward primer, 5’-TCACT GCAAT TGCCC ATGT-3’ (ACD6_HIF_Forward G-47729), targeting a conserved region of ACD6. PCR was performed using the program: 95°C for 2 min, 30 cycles of 95°C for 15 s, 55°C for 20 s, and 72°C for 45 s, followed by a final 5 min at 72°C and the products were separated on a 1% agarose gel. The amplification of only one primer pair was diagnostic of homozygous Est-1 or Col-0 sequences, while the amplification of both primer pairs indicated heterozygotes.

### 16S rRNA and ITS amplicon sequencing

Amplicons for microbial profiling were prepared as described in [34].

## Data availability

All sequencing data have been deposited in the European Nucleotide Archive (ENA, https://www.ebi.ac.uk/ena). They can be accessed under project number PRJEB78910.

## Author Contributions

DL and DW planned the study. DL and DW wrote the manuscript with input from all authors. SK, EMB, PPNA, KP, SS, and DM helped perform experiments. WY helped develop the RNAseq protocol. IB wrote image analysis software and assisted with photographic plant phenotyping. DL, SK, and KP collected samples and prepared libraries. WZ prepared genome edited Est:*acd6*-null lines. The “harvest team” comprises all the above authors plus: Adrián Contreras, Anjar Tri Wibowo, Bridgit Waithaka, Cristina Barragan, Efthymia Symeonidi, Frank Vogt, Hua Wang, Julia Elis, Lei Li, Moises Exposito Alonso, Or Shalev, Patricia Lang, Rui Wu, Sergio Latorre, Talia Karasov, Thanvi Srikant, and Ulrich Lutz.

## Acknowledgements

We thank Roosa Laitinen (University of Helsinki) for helpful comments on the manuscript and Eric Kemen (University of Tübingen) for helpful comments on experimental approaches. We thank Matt Horton and Jana Mittelstrass (University of Zürich) for helping to arrange planting facilities at Agroscope (Reckenholz) in Switzerland. The work has been supported by a Human Frontiers Science Program (HFSP) Long-Term Fellowship to DL (LT000565/2015-L), a Wallenberg Academy Fellows grant to DL (2021.0102), and ERC Advanced Grant IMMUNEMESIS 340602 and the Max Planck Society (DW).

## Competing Interests

DW holds equity in Computomics, which advises plant breeders. DW also consults for KWS SE, a globally active plant breeder and seed producer. The other authors declare no competing interests.

## Notes

### Competing Interest Statement

Detlef Weigel holds equity in Computomics, which advises plant breeders. Detlef Weigel also consults for KWS SE, a globally active plant breeder and seed producer. The other authors declare no competing interests.

