## Supplementary Material (PDF) for "Field conditions greatly modify a major growth-defense tradeoff in *Arabidopsis thaliana*"

### Supplementary Figure 1

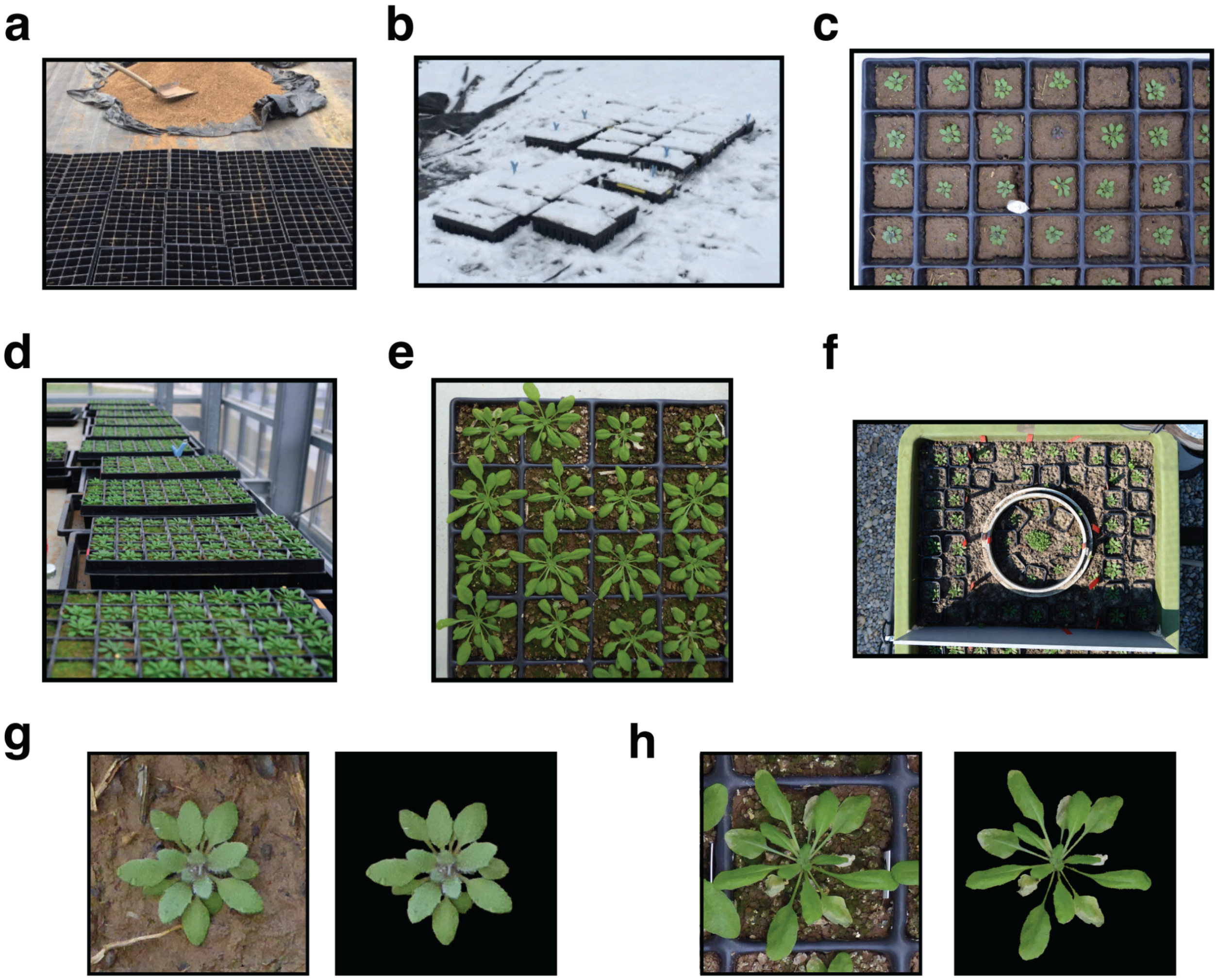


**Supplementary Fig. 1 | Plant cultivation. a,** Sieved and homogenized field soil ready for distribution into pots for experiments in Tübingen. **b,** Field-grown plants in Tübingen under snowfall in January. **c,** Field-grown plants in Tübingen from 2016-2017 season on the day of harvest. **d,** Greenhouse-grown plants in Tübingen from 2016-2017 season. **e,** Greenhouse-grown plants in Tübingen from 2016-2017 season on the day of harvest. **f,** Field-grown plants in Zurich from 2018-2019. **g,** Images of field plants from Tübingen on day or harvest before (left) and after (right) background removal. **h,** Images of greenhouse plants from Tübingen on day or harvest before (left) and after (right) background removal. Note that overlap of leaves from plants in adjacent pots occasionally required manual estimation of leaf borders.

### Supplementary Figure 2

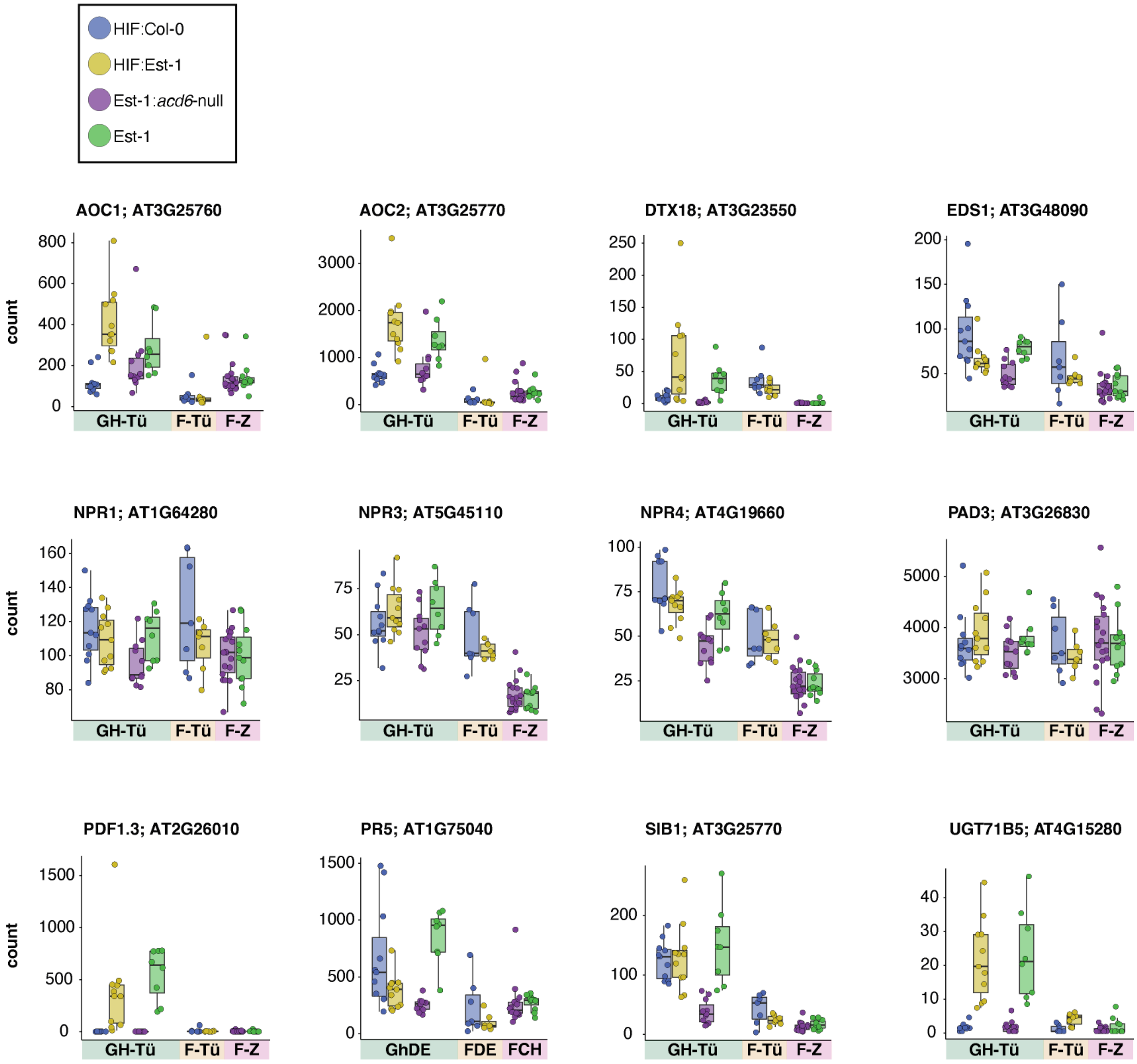


**Supplementary Fig. 2 | Expression of various immunity marker genes.** Normalized RNA-seq read counts (y-axes) for different genes with relevance to immune system activity. Colors of boxplots indicate plant genotype as described at the top left. The colored annotations along the x-axis denote the environment (GH-Tü = Greenhouse Tübingen, F-Tü = Field Tübingen, F-Z = Field Zurich).

### Supplementary Figure 3

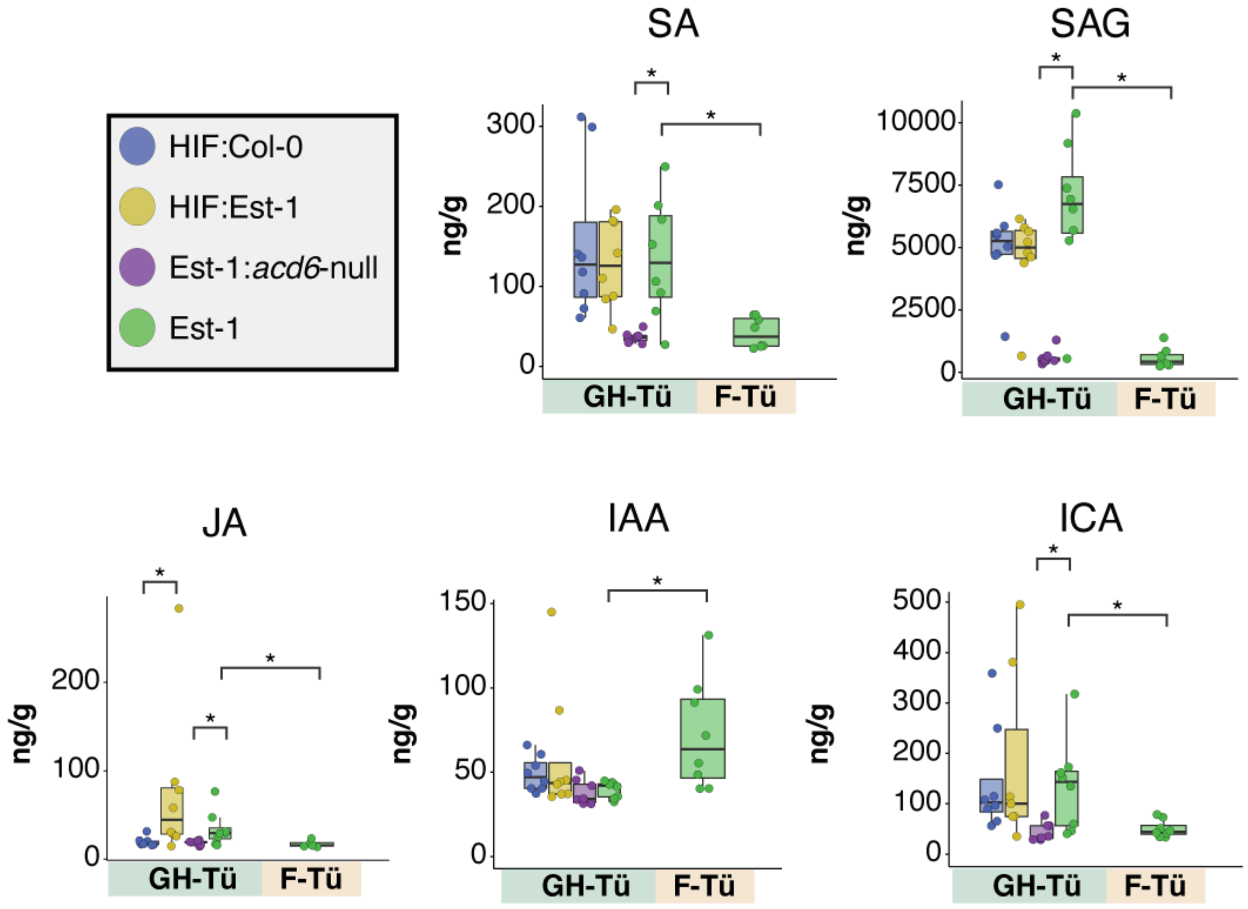


**Supplementary Fig. 3 | Phytohormone measurements.** From rosettes grown in Tübingen, salicylic acid (SA), its inactivate storage form SA O-β-glucoside (SAG), jasmonic acid (JA), indole-3-acetic acid (IAA), and indole-3-carboxylic acid (ICA) were measured from snap-frozen tissue via LC-MS. Letters “a” and “b” above the boxplots represent groups that are statistically different in a FDR-corrected Mann-Whitney U-test ($P < 0.05$).

### Supplementary Figure 4

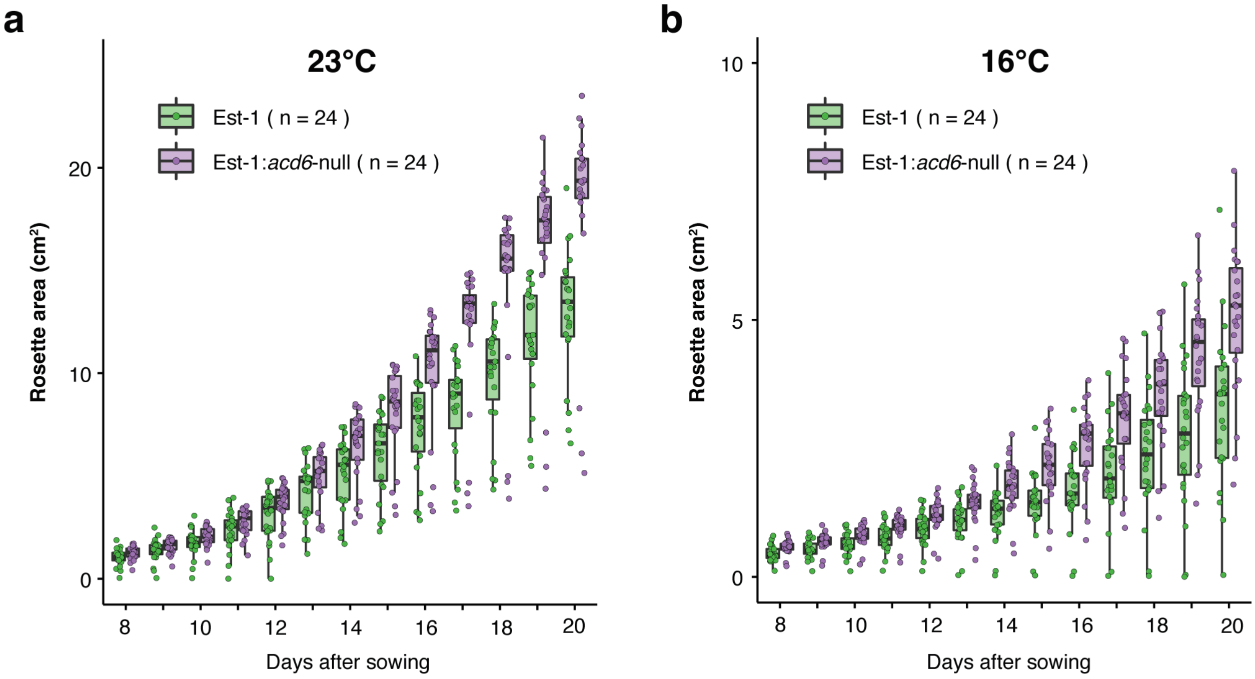


**Supplementary Fig. 4 | *ACD6*-dependent growth differences at 16°C and 23°C.** 24 Est-1 and 24 Est-1:*ACD6*-null seeds were sown in potting soil, grown in either 16°C or 23°C long (16 hr) days, and green pixels were monitored daily from overhead photographs starting 1 week after sowing. In both temperature regimes.

### Supplementary Figure 5

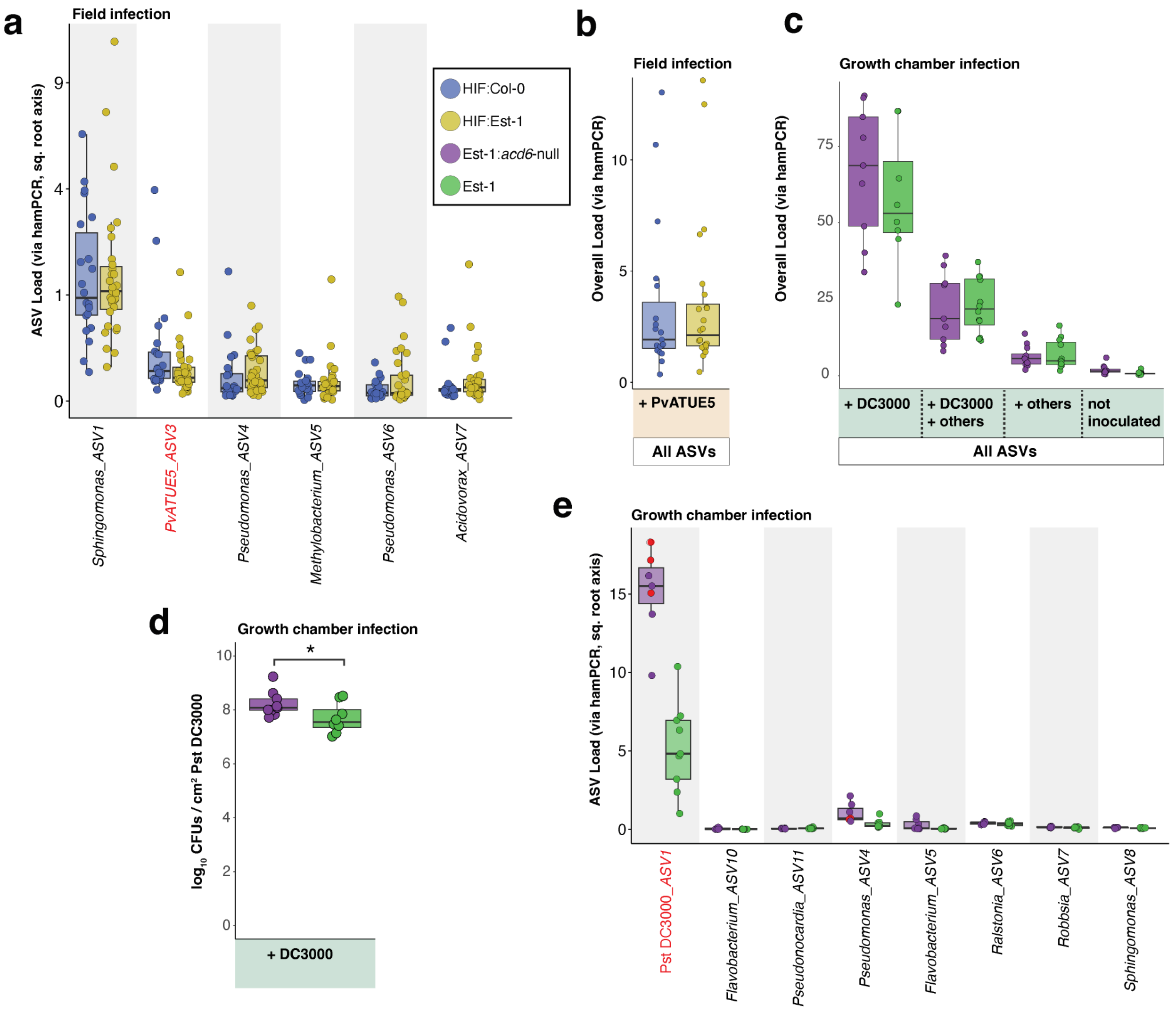


**Supplementary Fig. 5 | Artificial infections reveal subtle *ACD6*-dependent differences in bacterial assembly in the greenhouse. a,** Bacterial loads of the most abundant ASVs in field-grown HIF plants challenged with a cocktail of PvATUE5 strains in the field for 1 week, with color conventions for the plant genotypes as shown in the legend. The ASV corresponding to the cocktail of PvATUE5 is labeled in red. Boxes enclose the interquartile range (IQR) with whiskers extending to up to 1.5 times the IQR. **b,** Overall bacterial load considering all ASVs for the plants in (a). **c,** Overall bacterial load for growth chamber-grown Est-1 and Est-1:*acd6*-null plants challenged with Pst DC3000 and/or other phyllosphere bacteria for 4 days, with color code as in (a). **d,** Colony Forming Units (CFUs) of Pst DC3000 per cm$^2$ of leaf tissue for Pst DC3000-only infected plants shown in (c). * indicates $P < 0.05$ in a Mann-Whitney U-test. **e,** Bacterial loads of the most abundant ASVs in growth chamber-grown plants infected with Pst DC3000 only as shown in (c) and (d). Red points indicate samples with especially high bacterial load for which hamPCR could not provide accurate quantification due to a small number of reads from the plant's GIGANTEA gene. Sample size without considering the red points is too small and statistics are therefore omitted. The ASV corresponding to Pst DC3000 is labeled in red.
